# High-content screening identifies FDA-approved activators of mitophagy

**DOI:** 10.64898/2026.11.06.764995

**Authors:** Giacomo Giacchin, Valeria Balmaceda, Marta Giacomello, Andrea Eastes, Samantha Corrà, Lucía Fernández-Del-Rio, Nikole Scillitani, Davide Steffan, Marco Sandri, Erika Fernández-Vizarra, Orian Shirihai, Cristiane Benincá, Carlo Viscomi

**Author notes:** CONTACT: Carlo Viscomi, Department of Biosciences, University of Milano via Celoria 26-20133 Milano, Italy, and Veneto Institute of Molecular Institute (VIMM), via Orus 2-35129 Padova, Italy, OR, Cristiane Benincá, Department of Medicine, Division of Endocrinology David Geffen School of Medicine, University of California, 650 Charles E Young S Drive, CHS 27-100, 90095 Los Angeles, CA, USA.

## Abstract

Mitochondrial diseases are highly heterogeneous disorders caused by defects in oxidative phosphorylation, for which only limited therapeutic options are available. The elimination of dysfunctional mitochondria via mitophagy activation is an attractive strategy to treat several mitochondrial diseases, irrespective of the underlying genetic defect. To identify pharmacological activators of this pathway, we performed a high-content screening of 1,971 FDA-approved compounds using cellular reporters of TFEB nuclear translocation and mitophagy. This screening identified 294 compounds that promoted TFEB translocation, nine of which increased the number of mitolysosomes without compromising cell viability or damaging mitochondria. Among the selected compounds, Meptazinol and Ponatinib emerged as the top-ranked mitophagy inducers in fibroblasts from patients with complex I and IV deficiency, respectively, and were prioritised for further characterisation. However, they did not rescue the bioenergetic defect, suggesting that the stimulation of mitophagy per se is not sufficient to ameliorate the OXPHOS defect in mitochondrial diseases. Overall, our findings establish a scalable platform for the identification of pharmacological activators of TFEB and identify candidate compounds for therapeutic modulation of mitophagy.

**Summary blurb:** A two-step screen identifies approved drugs that boost the removal of damaged mitochondria without causing overt damage in both healthy cells and cells from patients with mitochondrial disease

## INTRODUCTION

Mitochondrial diseases are a group of genetically inherited conditions caused by an impairment of the oxidative phosphorylation (OXPHOS) system. Mutations in both nuclear and mitochondrial genes may cause dysfunction in one of the respiratory chain complexes (cI-IV), or adenosine triphosphate (ATP) synthase. Defects in cI and cIV are the most common cause of mitochondrial disease. No cure exists for these conditions and clinical management is limited to supportive therapy to relieve the symptoms (Gorman et al, 2016). Some therapeutic interventions have been tested and shown to be effective at the preclinical level, but only a few have moved to the clinical stage. For instance, the nicotinamide adenine dinucleotide precursor nicotinamide riboside, a component of vitamin B3, induced mitochondrial biogenesis by deacetylating and thus activating PPARG coactivator 1α, the master regulator of mitochondrial biogenesis (Cerutti et al, 2014; Khan et al, 2014). In a follow-up study, niacin, another vitamin B3 and adenine dinucleotide precursor, improved muscle strength in a small group of patients with progressive external ophthalmoplegia (Pirinen et al, 2020). Another promising preclinical approach for mitochondrial diseases exploited rapamycin, a well-known inhibitor of mammalian Target Of Rapamycin Complex 1 (mTORC1), a cytosolic Serine/Threonine kinase belonging to the phosphatidylinositol-4,5-bisphosphate 3-kinase-related protein kinases that control several cellular processes, including protein translation, autophagy, and immunity (Saxton & Sabatini, 2017). The beneficial effects of rapamycin have been demonstrated in several cellular and animal models of mitochondrial diseases (Johnson et al, 2013b; Zheng et al, 2016; Felici et al, 2017; Khan et al, 2017; Siegmund et al, 2017; Civiletto et al, 2018; Mito et al, 2022). Everolimus, a rapamycin analogue, was also shown to be effective in a few patients with mitochondrial encephalomyopathy with lactic acidosis and stroke-like episodes syndrome (MELAS) following kidney transplantation (Johnson et al, 2019) and in one patient with Leigh syndrome (Sage-Schwaede et al, 2019). However, rapamycin failed to ameliorate the phenotype of a mouse model of mitochondrial encephalopathy due to Coenzyme Q deficiency (Barriocanal-Casado et al, 2019), and in a clinical study in a child with MELAS syndrome (Sage-Schwaede et al, 2019).

We previously showed that rapamycin-dependent inhibition of mTORC1 promoted the coordinated activation of lysosomal biogenesis and autophagy via the activation of transcription factor EB (TFEB) (Civiletto et al, 2018). TFEB belongs to the microphthalmia/transcription factor E family of basic helix–loop–helix leucine zipper transcription factors and has been implicated in the transcriptional regulation of several cellular processes, such as autophagy and lysosomal and mitochondrial biogenesis (Settembre et al, 2011; Mansueto et al, 2017). TFEB is phosphorylated by mTORC1 and localizes to the cytosol (Settembre et al, 2012). Nutrient starvation or pharmacological inhibition of mTORC1 induce dephosphorylation and translocation of TFEB to the nucleus, where it triggers its transcriptional program (Sardiello et al, 2009). Acting downstream of mTORC1, TFEB is an ideal target for the development of new therapies for mitochondrial disorders aimed at enhancing the clearance of dysfunctional organelles. This can be achieved by activating mitochondria-specific autophagy (mitophagy), a cellular process aimed at selectively eliminating dysfunctional organelles. Here, we screened FDA-approved compounds to identify novel activators of mitophagy through a TFEB-based reporter assay and tested the top hit drugs in cellular models of mitochondrial disease.

## RESULTS

### Development of cellular models to investigate TFEB translocation and mitophagy

We first developed a reporter cell line to quantify mitophagy by isolating mouse adult fibroblasts (MAFs) from MitoQC-expressing transgenic mice (from now on MitoQC-MAFs) (McWilliams et al, 2016). The MitoQC reporter encodes a pH-sensitive fluorescent protein composed by tandem mCherry-GFP targeted to the outer mitochondrial membrane by the FIS1 leader peptide. When mitochondria undergo degradation and mitophagosomes fuse with lysosomes (mitolysosomes), the GFP signal is quickly quenched by the acidic environment, while the mCherry fluorescence remains visible. This allows the visualization of mitolysosomes as distinct red puncta and the quantification of mitophagy.

To validate our MitoQC-MAFs and identify a suitable positive control for mitophagy activation, we tested three concentrations of the reported autophagy/mitophagy activators rapamycin (Johnson et al, 2013a), Torin 1 (Thoreen et al, 2009), curcumin C1 (Song et al, 2016), and Urolithin A (UA) (Ryu et al, 2016). In basal conditions, we observed 6.4±1.0 mitolysosomes per cell, with a range between 0 and 19 puncta. We considered that a compound activated mitophagy when at least 20 mitolysosomes/cell were observed. Significant mitophagy induction was observed at 50 nM for Torin 1, 100 nM for rapamycin, 250 nM for curcumin C1, and 25 µM for UA (Figure 1A and B).

**Figure 1.**
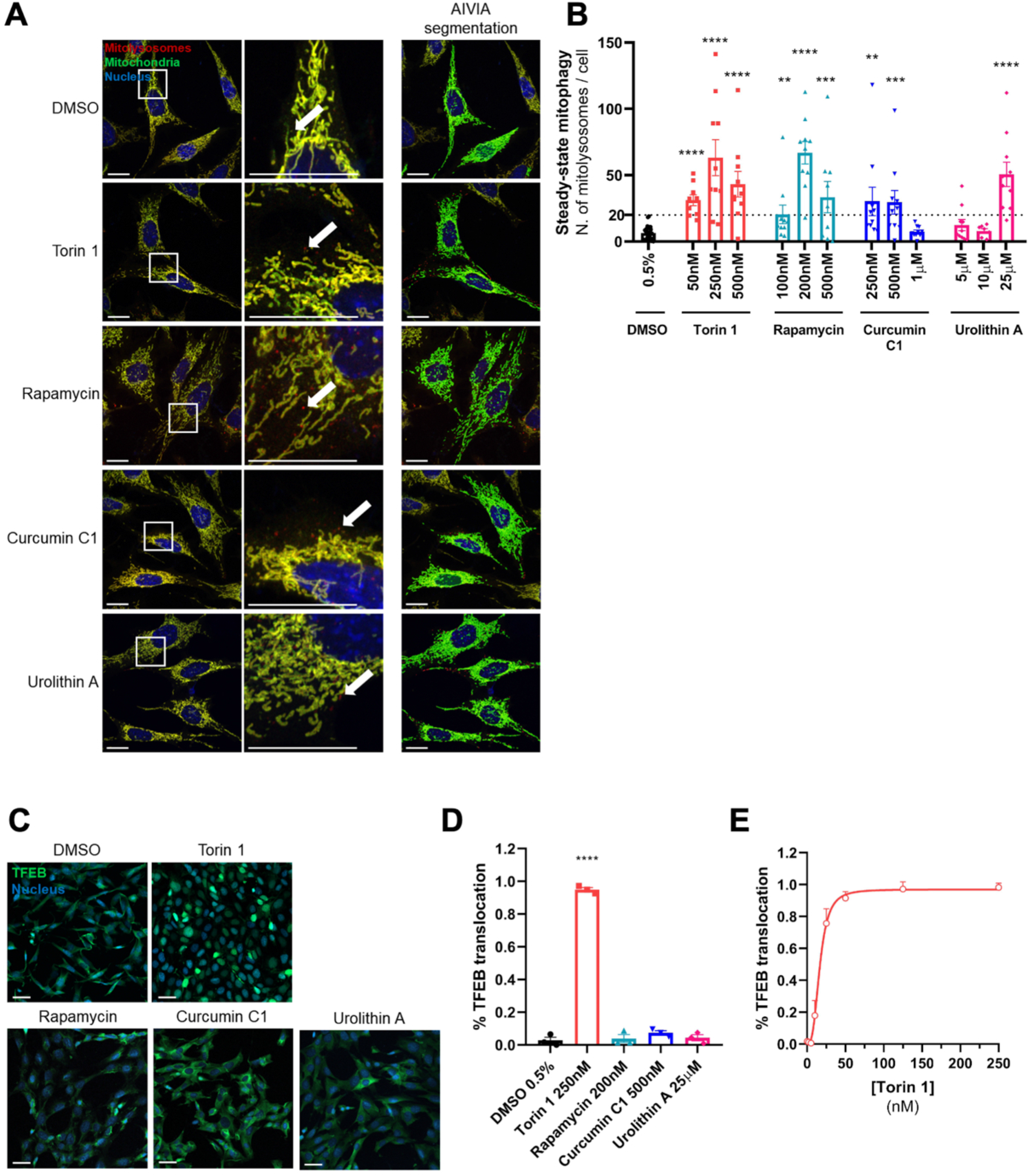
Validating cellular models and drug treatments to investigate mitophagy and TFEB translocation. (**A**) Representative Airyscan images (Z-stacks) of MitoQC-MAFs treated with 0.5% DMSO, Torin 1 50 nM, rapamycin 100 nM, curcumin C1 250 nM or Urolithin A 25 µM for 24 hours. Scale bars 20 µm. The arrows indicate mitolysosomes. (**B**) Quantification of steady-state mitophagy using AIVIA segmentation in MitoQC MAFs upon treatment with the mitophagy activators for 24 hours. N=6 biological replicates (DMSO), N=2 biological replicates (all other groups). A threshold for successful mitophagy activation is set at 20 mitolysosomes per cell. Data are expressed as mean ± SEM. Unpaired Student’s *t* tests were performed: Torin 1 vs DMSO all p<0.0001; rapamycin 100 nM vs DMSO p=0.0041; rapamycin 200 nM vs DMSO p<0.0001; rapamycin 500 nM vs DMSO p=0.0005; curcumin C1 250 nM vs DMSO p=0.0011; curcumin C1 500 nM vs DMSO p=0.0002; Urolithin A 25 µM vs DMSO p<0.0001. (**C**) Representative confocal images of TFEB-GFP MAFs treated with 0.5% DMSO, Torin 1 250 nM, rapamycin 200 nM, curcumin C1 500 nM or Urolithin A 25 µM for 24 hours. Scale bars 40 µm. (**D**) Quantification of TFEB translocation in TFEB-GFP MAFs upon treatment with mitophagy inducers for 24 hours. N=3 biological replicates. Data are expressed as mean ± SEM. One way ANOVA with Tukey’s post hoc multiple comparison test was performed: Torin 1 250 nM vs DMSO p<0.0001; all other comparisons were not significant. (**E**) Quantification of TFEB translocation in TFEB-GFP MAFs upon treatment with increasing concentrations (up to 250 nM) of Torin 1 for 24 hours. Non-linear fitting (variable slope, 4 parameters) for Torin 1 concentration vs. TFEB translocation: EC50=16.59 nM, R^2^=0.9862. N=5 biological replicates. Data are expressed as mean ± SEM.

To investigate whether the above compounds activated mitophagy via TFEB translocation to the nucleus, we generated TFEB-GFP over-expressing MAFs. Torin 1 induced robust TFEB translocation in 94.9±1.3% of the cells (p<0.0001), while rapamycin, curcumin C1, and UA had no effect at the concentrations used in our experiment (Figure 1C and D). Rapamycin lack of effect is not surprising, given that it acts as a weak inhibitor of mTORC1 in fibroblasts (Settembre et al, 2012). We then built a dose-response curve for Torin 1 up to 250 nM (Figure 1E). We observed that Torin 1 activated TFEB already at 10 nM (18.0±3.9% of TFEB-positive nuclei), with a half maximal effective concentration of 16.6 nM, and reached the plateau at 125 nM. Overall, these results indicate that Torin 1 is a powerful activator of both mitophagy and TFEB and we thus employed it as a positive control in the subsequent experiments at a saturating concentration of 125 nM.

### Screening FDA-approved drugs for the identification of TFEB-activating mitophagy inducers

We developed and optimised a workflow to identify new TFEB-activating compounds able to induce mitophagy, using confocal microscopy (Operetta CLS platform). We used Torin 1 treatment for 24 hours to validate our screening pipeline. We initially assessed TFEB translocation in MAFs over-expressing TFEB-GFP. Torin 1 increased TFEB-positive nuclei from 5.1±1.0% in untreated cells to 82.2±2.9% (p<0.0001) (Figure S1A and B). In addition, it markedly increased the number of mitolysosomes in MitoQC-MAFs from 8.8±1.0 to 57.0±6.4 per cell (p<0.0001) (Figure S1C and D). In the screening step we treated TFEB-GFP MAFs with a library of 1971 FDA-approved drugs for 24 hours (Figure 2A). Overall, a total of 294 compounds induced significant TFEB translocation compared to DMSO-treated cells (Figure 2B; Table S1). *In silico* analysis revealed that these compounds mostly impacted the cell cycle and DNA damage response, ubiquitination/proteasome, and proteases (Table S2). To account for toxicity, we scored the compounds based on cell count. Compounds falling below the 50^th^ percentile were considered toxic to our cells and excluded from subsequent experiments (Figure 2B; Table S1).

**Figure 2.**
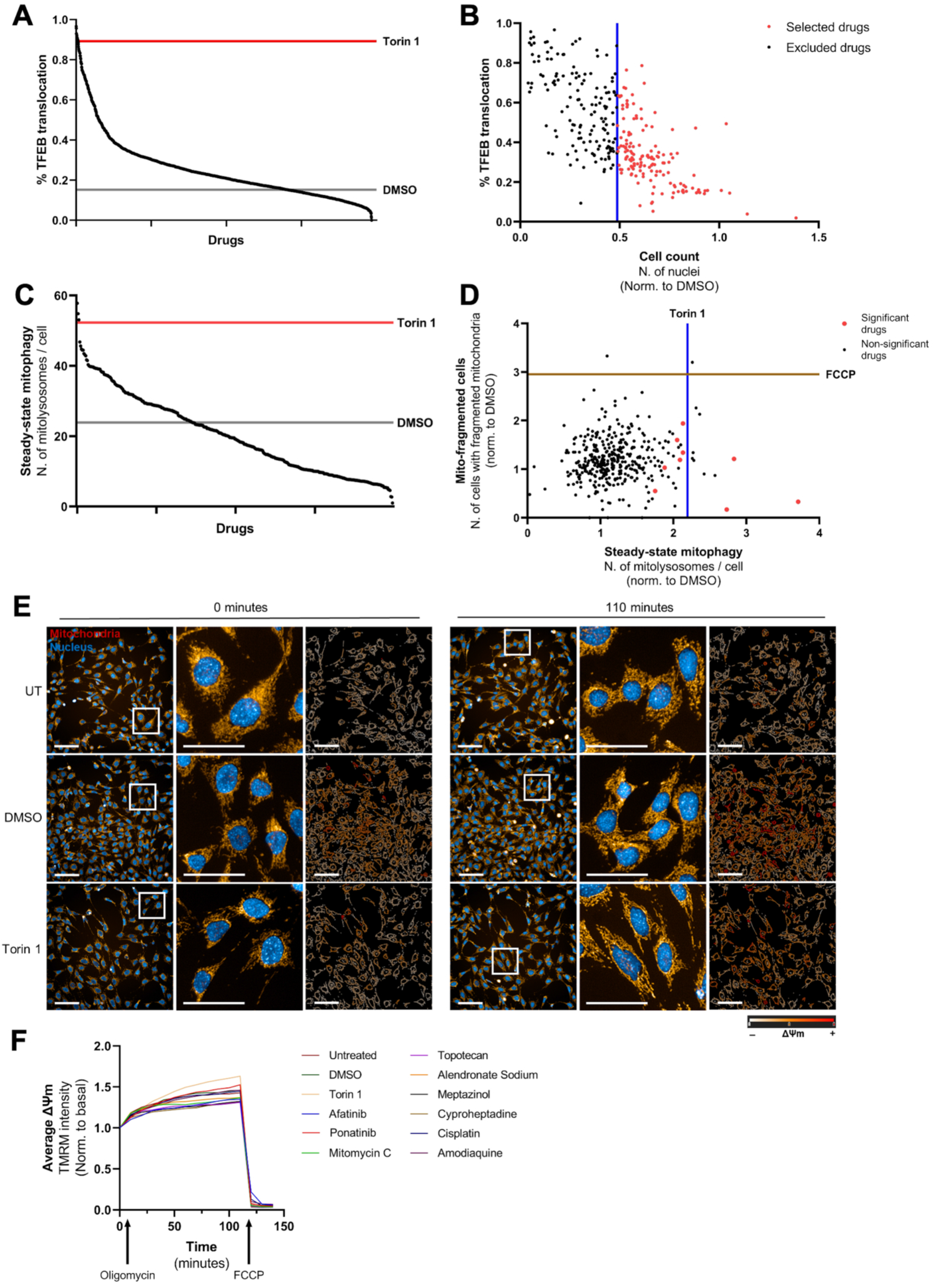
Screening FDA-approved drugs for TFEB-activators inducing mitophagy. (**A**) TFEB translocation ranked in descending order for each of the 1971 drugs, tested at 125 nM. Means of TFEB translocation for the negative and positive controls are indicated (15.2% for 1.25% DMSO, grey line; 89.3% for 125 nM Torin 1, red line). (**B**) Scatterplot of TFEB translocation over cell count for each of the 294 significant drugs. Drugs reducing cell count by >50% were excluded (black dots); drugs reducing cell count by <50% were selected for the MitoQC screening (red dots). The blue line indicates the threshold (cell count = 0.49 of DMSO). (**C**) Steady-state mitophagy ranked in descending order for each of the 147 drugs. Each drug was tested at three concentrations, ranging between 10 nM and 1 µM, for a total of 441 treatments. Means of mitolysosome number for the negative and positive controls are indicated (23.9 for 1.25% DMSO, grey line; 52.3 for 125 nM Torin 1, red line). (**D**) Scatterplot of mito-fragmented cells over steady-state mitophagy. Red dots indicate drugs significantly increasing mitophagy without mitochondrial fragmentation; black dots are non-significant drugs. Positive control Torin 1 at 125 nM (blue line) increased mitophagy by 2.2-folds vs DMSO-treated cells. Positive control FCCP at 5 µM (green line) increased the number of mito-fragmented cells by 2.9 folds vs DMSO-treated cells. (**E**) Representative fluorescence images of WT MAFs treated with 1.25% DMSO, 125 nM Torin 1, or 9 drugs at the selected concentration for 24 hours and stained with TMRM (red) and Hoechst (blue). Live cells were imaged in basal conditions (time = 0) and after addition of 1µM oligomycin and 5 µM FCCP (time = 140 minutes). Pseudocolored images represent changes in ΔΨm, measured as intensity of TMRM fluorescence. Scale bars 200 µm; zoomed images 100 µm. (**F**) Time-course of ΔΨm in untreated WT MAFs (black line) or treated with 1.25% DMSO (blue line), 125 nM Torin 1 (red line) or with the 9 selected drugs (grey lines) for 24 hours and upon addition of 1 µM oligomycin (time = 110 minutes) and 5 µM FCCP (time = 140 minutes).

The remaining 147 TFEB activators, which decreased cell count by less than 50%, were validated for their ability to induce mitophagy in MitoQC-MAFs (Figure 2C). Dose-response relationships for these compounds were also assessed at this validation step. For compounds above the 75^th^ percentile in cell count in the TFEB screening described above, we tested the following concentrations: 125 nM, 500 nM and 1 µM (Figure S1E). For the drugs below the 75^th^ percentile, we tested the following concentrations: 10 nM, 50 nM and 125 nM (Figure S1F). In addition, we simultaneously assessed number of mitolysosomes per cell and mitochondrial fragmentation, an index of mitochondrial health. Nine drugs produced a statistically significant increase in the number of mitolysosomes without fragmenting the mitochondrial network (Figure 2D) and were selected for the characterisation experiments at the concentrations detailed in Table S3.

To investigate whether the nine selected compounds influence mitochondrial bioenergetics, we used tetramethylrhodamine methyl ester perchlorate (TMRM) to analyse the mitochondrial membrane potential (ΔΨm) in basal conditions and after subsequent additions of oligomycin and FCCP in treated vs untreated cells. In “healthy” cells, oligomycin slightly hyperpolarizes ΔΨm by blocking the re-entry of protons through the ATP synthase. In contrast, cells with a compromised respiratory chain maintain their ΔΨm by reversing the activity of the same complex (Nicholls & Budd, 2000; Nicholls & Ward, 2000). We found that oligomycin hyperpolarised the ΔΨm in both untreated and treated cells (Figure 2E and F), indicating that the selected drugs had no uncoupling effect and thus do not induce mitophagy by damaging mitochondria.

### Characterization of mitophagy and TFEB activation in OXPHOS-defective human fibroblasts

We then tested the nine drugs selected from our screening in cellular models of mitochondrial disease. We stably expressed the MitoQC reporter in control and patient-derived fibroblasts harboring mutations in *NADH:ubiquinone oxidoreductase complex assembly factor 6* (*NDUFAF6*) (Catania et al, 2018) and in *SURF1 cytochrome c oxidase assembly factor* (*SURF1*) (Tiranti et al, 1998) (from now on named cI– and cIV-deficient cells, respectively). We then determined the mitophagy flux by calculating the ratio between the number of mitolysosomes per cell with and without inhibitors of the lysosomal hydrolases and proteases, Leupeptin and Pepstatin A. In cI-deficient cells, Meptazinol ranked as the top candidate among the nine drugs, showing the largest increase in mitophagy flux (38.5% over DMSO after 24 hours) (Figure 3A and C). Similarly, Ponatinib showed the largest increase in mitophagy flux in cIV-deficient fibroblasts (18.7%) (Figure 3B and D). Although this analysis for both compounds did not reach statistical significance in the multi-group comparison and given the exploratory nature of this additional characterisation step and the known variability of mitophagy flux measurements in patient-derived cells, we prioritised effect size, consistent with established approaches in high-throughput screening (Zhang et al, 2007). We thus selected Meptazinol and Ponatinib for further mechanistic and functional characterisation. Notably, both drugs also increased mitophagy flux in control cells by 25.2% (Meptazinol) and 13.0% (Ponatinib) (Figure S2A and B).

**Figure 3.**
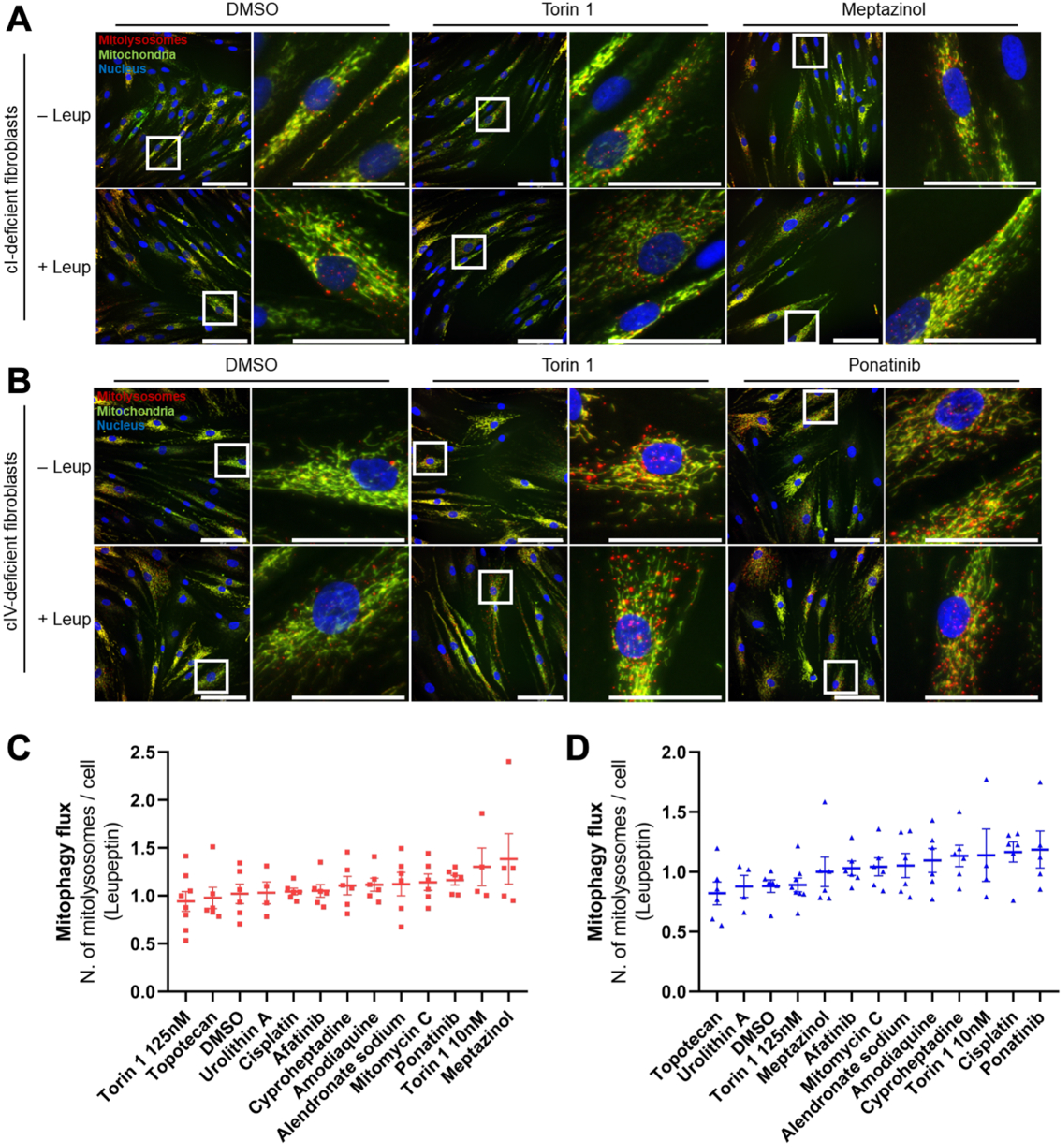
Mitophagy flux in OXPHOS-defective human fibroblasts treated with the 9 selected compounds. (**A**) Representative confocal images (single slices from Z-stacks) of MitoQC-expressing cI-deficient fibroblasts treated with 0.1% DMSO, 10 nM Torin 1 or 50 nM Meptazinol for 24 hours, plus co-treatment with the lysosomal inhibitors (Leupeptin). Nuclei are stained with DAPI (blue). For each treatment low magnification images (scale bars 100 µm) and zoomed images in the white square (scale bars 50 µm) are shown. (**B**) Representative confocal images (single slices from Z-stacks) of MitoQC-expressing cIV-deficient fibroblasts treated with 0.1% DMSO, 10 nM Torin 1 or 50 nM Ponatinib for 24 hours, plus co-treatment with the lysosomal inhibitors (Leup). Nuclei are stained with DAPI (blue). Scale bars 100 µm, zoomed images 50 µm. (**C**) Quantification of mitophagy flux in MitoQC-expressing cI-deficient fibroblasts upon treatment with 9 selected drugs, Torin 1 (10 nM and 125 nM), Urolithin A (50 µM) or DMSO (0.1%) for 24 hours. Data are expressed as mean ± SEM, ranked from least to most effective drug. N=4 biological replicates (Urolithin A, Torin 1 10 nM); N=5 biological replicates (Meptazinol); N=8 biological replicates (Torin 1 125 nM); N=6 biological replicates (all other groups). One way ANOVA with Tukey’s post hoc multiple comparison test was performed: all comparisons were not significant. (**D**) Quantification of mitophagy flux in MitoQC-expressing cIV-deficient fibroblasts upon treatment with 9 selected drugs, Torin 1 (10 nM and 125 nM), Urolithin A (50 µM) or DMSO (0.1%) for 24 hours. Data are expressed as mean ± SEM, ranked from least to most effective drug. N=4 biological replicates (Urolithin A, Torin 1 10 nM); N=5 (Ponatinib); N=8 (Torin 1 125 nM); N=6 biological replicates (all other groups). One way ANOVA with Tukey’s post hoc multiple comparison test was performed: all comparisons were not significant.

We then validated TFEB engagement by Meptazinol and Ponatinib in cI-and cIV-deficient fibroblasts, respectively. Here, we assessed endogenous TFEB activation by immunofluorescence in human-derived cells, whereas the primary screen relied on TFEB-overexpressing mouse fibroblasts. Both cI (5.108±1.163%, p<0.0001) and cIV-deficient (3.843±1.094%, p<0.0001) fibroblasts exhibited a significantly reduced proportion of TFEB-positive nuclei compared to control fibroblasts (22.530±1.495%) (Figure S3A and B). Treatment with Torin 1 induced rapid TFEB nuclear translocation within 3 hours in control (45.130±6.815%, p=0.0004) (Figure 4A and B), cI-deficient (17.550±3.182%, p=0.0002) (Figure S3C and D) and cIV-deficient fibroblasts (29.620±8.888%, p=0.0140) (Figure 4D and E), confirming pathway responsiveness. TFEB nuclear localisation returned to baseline levels after 16 hours of treatment with Torin 1 in mutant cells (Figure 4D and F, Figure S3C and E), but not in control fibroblasts (36.53±6.224%, p=0.0441) (Figure 4A and C). Ponatinib induced a delayed but sustained TFEB nuclear translocation after 16 hours of treatment in control (42.54±1.541%, p=0.0002) (Figure 4A and C) and cIV-deficient cells (11.130±2.758%, p=0.0278) (Figure 4D and F). Instead, Meptazinol promoted TFEB nuclear translocation in control (31.62±1.868%, p=0.0132) (Figure 4A-C), but not in cI-deficient fibroblasts (Figure S3C-E).

**Figure 4.**
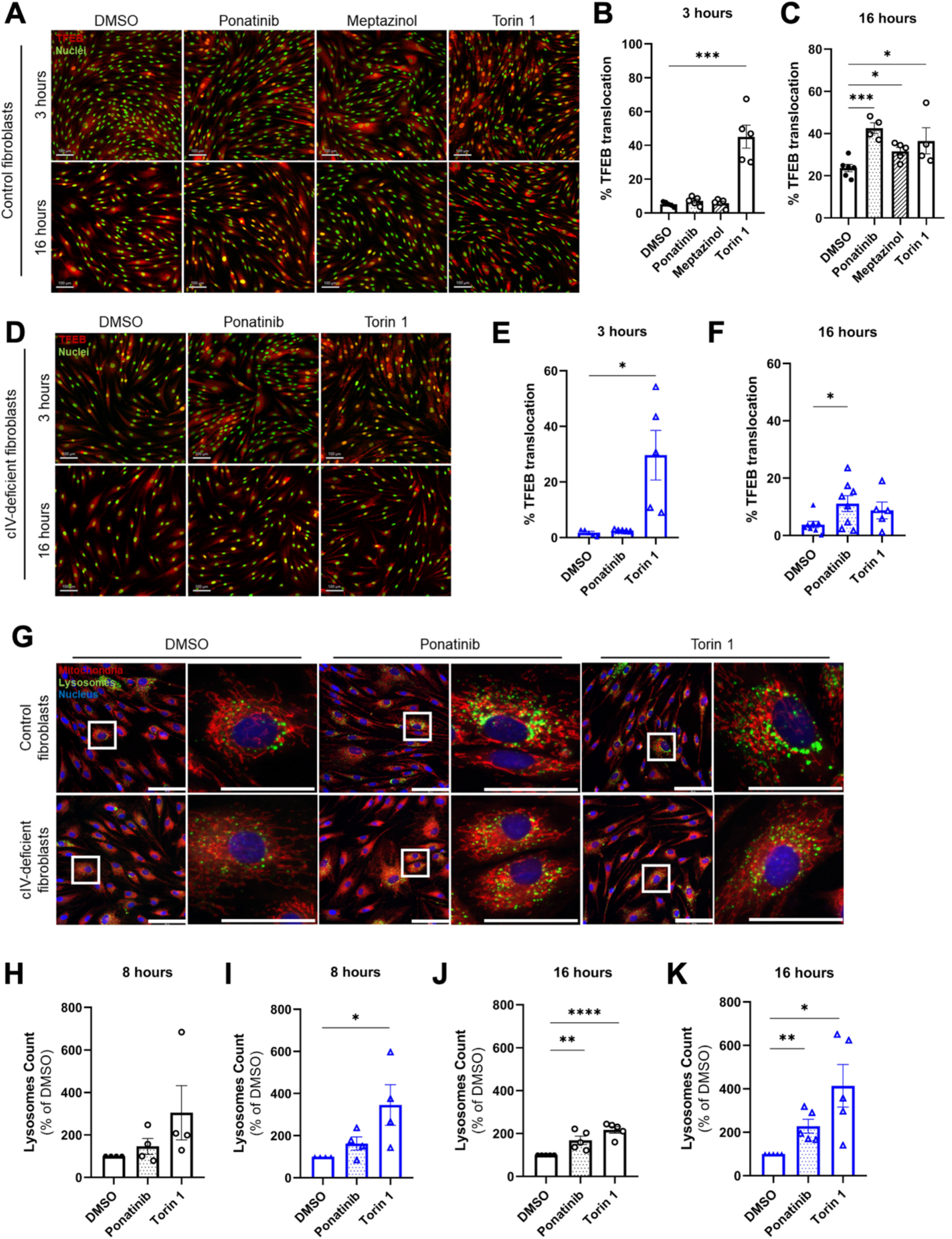
Ponatinib induces TFEB translocation to the nucleus and lysosomal biogenesis in control and cIV-deficient human fibroblasts. (**A**) Representative confocal images of control fibroblasts treated for 3 or 16 hours with 50 nM Ponatinib, 50 nM Meptazinol, or 125 nM Torin 1 and stained with TFEB (red). Nuclei are stained with DAPI (green). Scale bars 100 µm. (**B**) Quantification of the percentage of TFEB positive nuclei in control fibroblasts following a 3-hour treatment with 50 nM Ponatinib, 50 nM Meptazinol, or 125 nM Torin 1. Unpaired Student’s *t* tests were performed: Torin 1 vs DMSO p=0.0004. (**C**) Quantification of the percentage of TFEB positive nuclei in control fibroblasts following 16 hours treatment with 50 nM Ponatinib, 50 nM Meptazinol, or 125 nM Torin 1. Unpaired Student’s *t* tests were performed: Ponatinib vs DMSO p=0.0002; Meptazinol vs DMSO p=0.0132; Torin 1 vs DMSO p=0.0441. (**D**) Representative confocal images of cIV-deficient fibroblasts treated for 3 or 16 hours with 50 nM Ponatinib or 125 nM Torin 1 and stained with TFEB (red). Nuclei are stained with DAPI (green). Scale bars 100 µm. (**E**) Quantification of the percentage of TFEB positive nuclei in cIV-deficient fibroblasts following 3 hours and (**F**) 16 hours of treatment with 50 nM Ponatinib or 125 nM Torin 1. Unpaired Student’s *t* tests were performed: Torin 1 vs DMSO (3 hours) p=0.0140; Ponatinib vs DMSO (16 hours) p=0.0278; all other p-values were not significant. (**G**) Representative confocal images of control and cIV-deficient fibroblasts treated for 16 hours with 50 nM Ponatinib or 125 nM Torin 1. Mitochondrial are stained with MitoTracker (red), lysosomes with LysoTracker (green), and nuclei with Hoechst (blue). Scale bars 100 µm, zoomed images 50 µm. (**H**) Quantification of lysosome number per cell in control and (**I**) cIV-deficient fibroblasts following 8 hours treatment with 50 nM Ponatinib or 125 nM Torin 1. Lysosome count was normalized to the DMSO condition. Unpaired Student’s *t* tests were performed: cIV-deficient fibroblasts Torin 1 vs DMSO p=0.0436; all other p-values were not significant. (**J**) Quantification of total number of lysosomes per cell in control fibroblasts and (**K**) cIV-deficient fibroblasts following 16 hours of treatment with 50 nM Ponatinib or 125 nM Torin 1. Lysosome count was normalized to the DMSO condition. Unpaired Student’s *t* tests were performed: control fibroblasts Ponatinib vs DMSO p=0.0076; Torin 1 vs DMSO p<0.0001; cIV-deficient fibroblasts Ponatinib vs DMSO p=0.0039; Torin 1 vs DMSO p=0.0125. For all experiments N ≥ 4 biological replicates.

To further define the mechanism of Ponatinib– and Meptazinol-mediated TFEB activation in control fibroblasts, we examined the phosphorylation levels of ribosomal protein S6, a downstream target mTORC1. As expected, Torin 1 reduced S6 phosphorylation (0.2435±0.0186, p<0.0001). Ponatinib similarly decreased S6 phosphorylation (0.6258±0.0899, p=0.0021), although to a lesser extent than Torin 1 (Figure S3F and G), indicating that Ponatinib activates TFEB via mTORC1 inhibition. In contrast, Meptazinol treatment did not alter S6 phosphorylation levels in control fibroblasts (0.8910±0.1039), suggesting that its effect on TFEB is independent of mTORC1 signalling (Figure S3F and G).

Since Ponatinib activated TFEB in both control and cIV fibroblasts, we analysed lysosomal content by performing LysoTracker staining. Lysosome count increased upon Torin 1 treatment as early as 8 hours in cIV-deficient cells (345.6±94.6, p=0.0436) but not in control cells (304.4±127.7) (Figure 4H and I), supporting a rapid TFEB activation for Torin 1. Moreover, Torin 1 increased lysosome count in both cell types at 16 hours (217.0±15.3 in control cells, p<0.0001; 413.7±97.9 in cIV-deficient cells, p=0.0125) (Figure 4G, J and K). Ponatinib showed no early changes in lysosomal content at 8 hours (Figure 4H and I). However, after16 hours of treatment, Ponatinib increased lysosome counts in both control (168.0±19.2, p=0.0076) and cIV-deficient cells (227.9±31.9, p=0.0039) (Figure 4G, J and K), consistent with the delayed TFEB activation observed. Together with mTORC1 inhibition and TFEB nuclear translocation, these data indicate that Ponatinib promotes remodelling of the autophagy-lysosome pathway, creating a cellular environment permissive for improved mitochondrial quality control.

### cI– and cIV-defective patient fibroblasts accumulate dysfunctional mitochondria and have reduced mitochondrial respiration

When mitochondria are damaged or dysfunctional, their ΔΨm is reduced, they fragment and undergo mitophagy (Twig & Shirihai, 2011). We investigated mitochondrial ΔΨm and morphology by staining our cells with both tetramethylrhodamine ethyl ester perchlorate (TMRE) and MitoTracker Green (MTG). The average intensity of mitochondrial TMRE was similar in cI-, cIV-deficient, and control fibroblasts, indicating that overall ΔΨm is unaffected in the mutant fibroblasts (Figure S4A and B). However, we observed an increase in the number of small mitochondria per cell in cI-deficient (30.5±1.5, p=0.0007) and cIV-deficient cells (28.6±1.0, p=0.0002) compared to control fibroblasts (20.7±0.2) (Figure S4C). Notably, a greater proportion of these small mitochondria displayed reduced ΔΨm in both cI-deficient (19.5±0.5, p<0.0001) and cIV-deficient cells (15.5 ± 2.1, p=0.0290) relative to controls (9.4 ± 0.6) (Figure S4D). These data indicate that fibroblasts with defects in cI and cIV accumulate partially fragmented and depolarized mitochondria before targeting them to mitophagy.

We then evaluated how defects in cI and cIV affected energy metabolism of patient-derived fibroblasts by using Seahorse assay. In basal conditions, cI-deficient cells had a significantly reduced oxygen consumption rate (OCR) (21%, p=0.0005) and OXPHOS-related ATP production (21%, p=0.0012) (Figure S4E and G). We also observed a mild reduction in basal (14%, p=0.002) and oligomycin-induced (30%, p<0.0001) extracellular acidification rate (ECAR) in cI-deficient vs control fibroblasts, leading to a 12% reduction in glycolysis-related ATP synthesis (p=0.0114) (Figure S4F and G). This finding may reflect accumulation of NADH in cI-deficient cells, with a consequent reduction in NAD⁺ availability, an essential cofactor for glycolysis, potentially limiting flux through GAPDH (Xie et al, 2020). In contrast, cIV-deficient cells showed a significant reduction in OCR (48%, p<0.0001) and OXPHOS-dependent ATP production rate (46%, p<0.0001) compared to control fibroblasts (Figure S4E and G). Basal ECAR, oligomycin-induced ECAR and glycolysis-related ATP synthesis significantly increased by 82% (p<0.0001), 32% (p=0.0048), and 101% (p<0.0001), respectively, in cIV-deficient fibroblasts compared to the control cells (Figure S4F and G).

Overall, these data confirm that defects in cI and cIV, caused by mutations in *NDUFAF6* and *SURF1* respectively, contribute to a reduced mitochondrial respiration and OXPHOS-related ATP production rates in patient-derived fibroblasts. In addition, they also indicate that the rather severe cIV defect rewires metabolism toward glycolysis. This is not observed in cI-defective cells, which have a milder defect.

### Meptazinol and Ponatinib promote mitochondrial fragmentation and mitophagy without restoring respiratory capacity in OXPHOS-deficient fibroblasts

We first tested whether Meptazinol affects ΔΨm. Meptazinol did not depolarise the mitochondrial membrane in control or cI-deficient fibroblasts (Figure S5A, B, E, F). In control cells, it increased the number of small (30.1±3.3, p=0.0278) and small-depolarised mitochondria (13.2±1.1, p=0.0242) without causing generalised fragmentation or depolarisation of the network (Figure S5C and D), whereas neither parameter changed in cI-deficient cells (Figure S5G and H).

We next assessed energy metabolism by Seahorse assay. Meptazinol left OCR, ECAR and both OXPHOS– and glycolysis-derived ATP production unchanged in control (Figure S6A-C) and cI-deficient fibroblasts (Figure S6D-F). Prolonging treatment to 72 hours did not alter basal or maximal oxygen flux in permeabilised cells measured by high-resolution respirometry (Oroboros) in either genotype (Figure S6G and H). Meptazinol therefore induces mitophagy without affecting membrane potential or ATP production.

We performed the same analysis in Ponatinib-treated cIV-deficient fibroblasts. Ponatinib did not reduce average ΔΨm in cIV-deficient or control cells (Figure 5A and B; Figure S5A and B) but increased small mitochondria from 28.6±1.0 to 40.8±2.4 in cIV-deficient cells (p=0.0034; Figure 5C) and from 20.7±0.2 to 26.8±1.9 in controls (p=0.0179; Figure S5C). Small-depolarised mitochondria rose from 15.5±2.1 to 22.9±1.1 in *SURF1*-mutant cells (p=0.0189; Figure 5D) and from 9.4±0.6 to 14.7±1.4 in controls (p=0.0115; Figure S5D), consistent with segregation of damaged mitochondria for mitophagic clearance.

**Figure 5.**
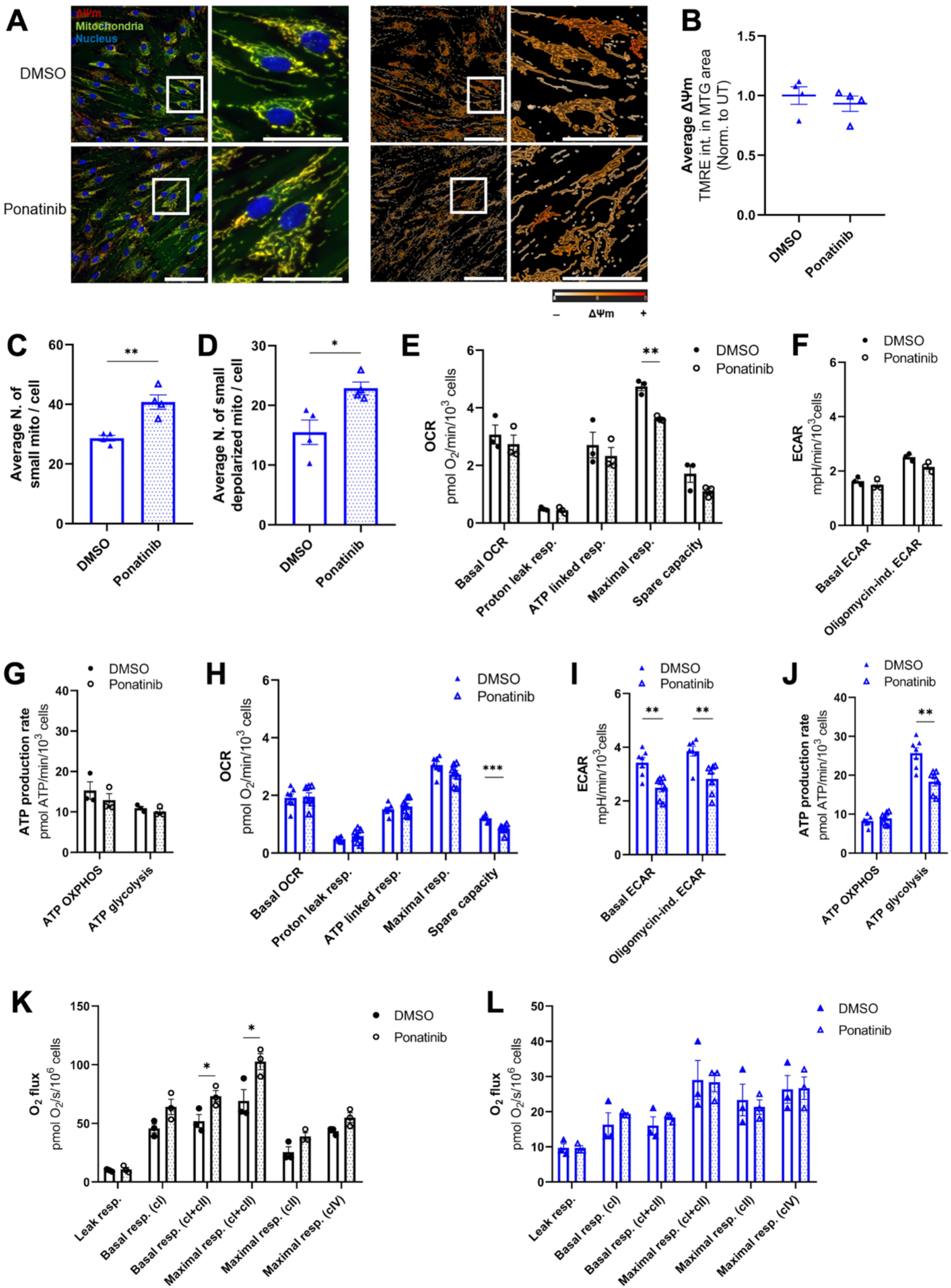
Ponatinib fragments mitochondria and reduces glycolysis-linked ATP production in cIV-defective fibroblasts. (**A**) Representative confocal images (single slices from Z-stacks) of cIV-deficient fibroblasts treated with 0.1% DMSO or 50 nM Ponatinib for 24 hours and stained with TMRE (red) plus MitoTracker Green (green). Nuclei are stained with Hoechst (blue). Pseudocolored images represent changes in ΔΨm, measured as intensity of TMRE fluorescence. Scale bars 100 µm, zoomed images 50 µm. (**B**) Quantification of average ΔΨm per cell in cIV-deficient fibroblasts treated with 0.1% DMSO or 50 nM Ponatinib, normalized to average signal in DMSO condition. N=4 biological replicates. Data are expressed as mean ± SEM. An unpaired Student’s *t* test was performed: p-value not significant. (**C**) Quantification of average number of small mitochondria (area <1 µm^2^) per cell in cIV-deficient fibroblasts treated with 0.1% DMSO or 50 nM Ponatinib. N=4 biological replicates. Data are expressed as mean ± SEM. An unpaired Student’s *t* test was performed: Ponatinib vs DMSO p=0.0034. (**D**) Quantification of average number of small-depolarized mitochondria per cell in cIV-deficient fibroblasts treated with 0.1% DMSO or 50 nM Ponatinib. N=4 biological replicates. Data are expressed as mean ± SEM. An unpaired Student’s *t* test was performed: Ponatinib vs DMSO p=0.0189. (**E**) Oxygen Consumption Rates in control fibroblasts, after a 24-hour treatment with 0.1% DMSO or 50 nM Ponatinib. Data are expressed as mean ± SEM. N=3 biological replicates. Unpaired Student’s t tests were performed: maximal respiration Ponatinib vs DMSO p=0.0018; all other p-values were not significant. (**F**) Basal Extracellular Acidification Rates in control fibroblasts, after a 24-hour treatment with 0.1% DMSO or 50 nM Ponatinib. Data are expressed as mean ± SEM. N=3 biological replicates. Unpaired Student’s t tests were performed: all p-values were not significant. (**G**) ATP production rates in control fibroblasts, after a 24-hour treatment with 0.1% DMSO or 50 nM Ponatinib. Data are expressed as mean ± SEM. N=3 biological replicates. Unpaired Student’s t tests were performed: all p-values were not significant. (**H**) Oxygen Consumption Rates in cIV-deficient fibroblasts, after a 24-hour treatment with 0.1% DMSO or 50 nM Ponatinib. Data are expressed as mean ± SEM. N=7 biological replicates. Unpaired Student’s *t* tests were performed: spare capacity Ponatinib vs DMSO p=0.0004; all other p-values were not significant. (**I**) Basal Extracellular Acidification Rates in cIV-deficient fibroblasts, after a 24-hour treatment with 0.1% DMSO or 50 nM Ponatinib. Data are expressed as mean ± SEM. N=7 biological replicates. Unpaired Student’s *t* tests were performed: basal ECAR Ponatinib vs DMSO p=0.0021; oligomycin-induced ECAR Ponatinib vs DMSO p=0.0029. (**J**) ATP production rates in cIV-deficient fibroblasts, after a 24-hour treatment with 0.1% DMSO or 50 nM Ponatinib. Data are expressed as mean ± SEM. N=7 biological replicates. Unpaired Student’s *t* tests were performed: ATP from glycolysis Ponatinib vs DMSO p=0.0014; all other p-values were not significant. (**K**) Oxygen flux per cell in permeabilized cIV-deficient fibroblasts, after a 72-hour treatment with 0.1% DMSO or 50 nM Ponatinib. Data are expressed as mean ± SEM. N=3 biological replicates. Unpaired Student’s *t* tests were performed: all p-values were not significant. (**L**) Oxygen flux per cell in permeabilized control fibroblasts, after a 72-hour treatment with 0.1% DMSO or 50 nM Ponatinib. Data are expressed as mean ± SEM. N=3 biological replicates. Unpaired Student’s t tests were performed: basal respiration (cI+cII) Ponatinib vs DMSO p=0.0466; maximal respiration (cI+cII) Ponatinib vs DMSO p=0.0468; all other p-values were not significant.

Ponatinib lowered maximal OCR (p=0.0018) without affecting ATP-linked respiration, OXPHOS-linked ATP production (Figure 5E and G), or basal and oligomycin-induced ECAR in control cells (Figure 5F). In cIV-deficient fibroblasts, it reduced maximal respiration (p=0.0945) and spare capacity (p=0.0004) (Figure 5H) and lowered basal and oligomycin-induced ECAR by 27% (p=0.0021 and p=0.0029; Figure 5I), cutting glycolytic ATP production by 40% (p=0.0014; Figure 5J). At 72 hours of treatment, Ponatinib increased basal (51.95 to 73.10 pmol O₂/s, p=0.0466) and maximal respiration (69.15 to 102.60 pmol O₂/s, p=0.0468) in permeabilized control fibroblasts (Figure 5K), suggesting that the decrease seen at 24 hours was transient. Oxygen flux in permeabilised cIV-deficient cells was unchanged (Figure 5L).

Together, these data show that Ponatinib promotes the segregation and clearance of depolarised mitochondria in cIV-deficient fibroblasts, but that enhanced quality control alone does not restore respiratory function within the timeframe tested.

## DISCUSSION

In this work, we established a scalable screening platform to identify pharmacological compounds that promote TFEB activation and enhance mitophagy. Using this pipeline, we identified and validated two FDA-approved compounds able to modulate mitochondrial quality control, while also revealing that enhanced quality control alone is not sufficient to restore respiratory function in cellular models of OXPHOS dysfunction. Our screening identified 294 pharmacological TFEB activators. This large number of initial hits reflects the central role of TFEB in cellular adaptation to diverse cellular stressors, as multiple signalling pathways converge on its regulation (Takla et al, 2023). Therefore, the validation step was crucial in narrowing down the hit compounds to those specifically activating mitophagy. A key innovation in this step is the simultaneous assessment of mitochondrial toxicity and mitophagy by assessing mitochondrial fragmentation and mitolysosome number, which allowed us to exclude compounds that induce mitophagy primarily as a consequence of generalised mitochondrial damage (Allen et al, 2013). This approach led to the identification of nine mitophagy activators.

As mitophagy is a tightly regulated and dynamic process, even moderate increases in mitophagy flux may reflect meaningful changes in mitochondrial quality control. However, whether these changes translate into improved mitochondrial function depends on the balance between mitochondrial clearance, replacement, and underlying bioenergetic defect. Conversely, an excessive increase in mitophagy flux can be detrimental, particularly in already vulnerable mitochondrial disease contexts (Borbolis & Palikaras, 2022; Gao et al, 2024). Accordingly, our selection criteria prioritised nine compounds that significantly increased steady-state number of mitolysosomes without inducing cellular or mitochondrial toxicity, followed by their validation of enhanced mitophagy flux in patient-derived cells.

Among these, Meptazinol and Ponatinib showed the greatest increases in mitophagy flux in cI– and cIV-deficient patients’ fibroblasts and were selected for further mechanistic characterisation. Meptazinol induced mitophagy independently of mTORC1 signaling, whereas Ponatinib promoted mTORC1-dependent TFEB activation accompanied by increased lysosomal biogenesis. These findings provide independent evidence that both compounds modulate mitochondrial quality control pathways, albeit through distinct mechanisms. Notably, although Meptazinol was originally identified in our screening based on its ability to induce TFEB nuclear translocation in TFEB-overexpressing mouse fibroblasts and was confirmed in human control cells, it failed to promote endogenous TFEB activation in cI-deficient cells. This observation underscores the importance of the multi-tiered validation strategy employed in this study, which combined phenotypic screening with functional and mechanistic characterization in patient-derived cellular models.

Meptazinol is a synthetic analgesic with partial µ-opioid receptor agonist activity and a relatively good safety profile with low abuse potential and tolerance, and only minor respiratory depressive effects (Franklin & Aldridge, 1976; Spiegel & Pasternak, 1984; Holmes & Ward, 1985). Its mild activity on the μ-opioid receptors explains why Meptazinol is only one-tenth as potent as morphine (Hoskin & Hanks, 1991; Pasternak & Pan, 2013). Moreover, Meptazinol inhibits acetylcholinesterase (AChE) (Strahan et al, 1985; Galli et al, 1996), which are highly expressed but with low catalytic activity in fibroblasts, where it localises to the leading edge of migrating cells during wound healing (Anderson et al, 2008) (Bigliardi-Qi & Bigliardi, 2015). While opioids can regulate autophagy, no link has been established between AChE activity and autophagy. Morphine activates Beclin 1– and ATG5-dependent autophagy in neuronal cells (Zhao et al, 2010), and k-opioid receptor stimulation induces autophagy though ERK1/2-dependent upregulation of LC3-II, Beclin 1 and ATG5 (Karoussiotis et al, 2022). As for other opioids, Meptazinol may therefore induce autophagy, which warrants future studies. Notably, Meptazinol increased the number of small and small-depolarized mitochondria in control cells and enhanced mitophagy flux in cI-deficient cells without affecting respiration, glycolysis nor ATP production, indicating that mitophagy induction alone was insufficient to restore mitochondrial function.

Ponatinib is a multi-target tyrosine kinase inhibitor (Tan et al, 2019) approved for chronic myeloid leukemia and Philadelphia chromosome-positive acute lymphoblastic leukemia (Cortes et al, 2012). In addition, it has shown efficacy in other conditions, including pulmonary arterial hypertension and pulmonary fibrosis, by suppressing the TGF-β1/Smad3 pathway (Qu et al, 2015; Kang et al, 2016), as well as cerebral cavernous malformation through MEKK3 kinase inhibition (Choi et al, 2018). Since TGF-β/Smad3 signalling predominantly activates mTORC1 in many contexts (Das et al, 2013; Rozen-Zvi et al, 2013), inhibition of this pathway represents one potential mechanism through which Ponatinib could modulate mTORC1 signalling and promote TFEB activation, although this relationship remains to be established in our experimental models. Recently, Ponatinib was reported to restore autophagy in non-alcoholic steatohepatitis through the normalization of TFEB transcript and its target genes (Lin et al, 2025). Here, we show that Ponatinib induces TFEB nuclear translocation and mitophagy through an mTORC1-dependent mechanism in human fibroblasts. In addition, Ponatinib treatment increased the number of small and small-depolarized mitochondria in cIV-deficient cells. One possible interpretation is that Ponatinib stimulates peripheral fission, a process that precedes the removal of damaged mitochondria via mitophagy and that requires local ΔΨm depolarization (Kleele et al, 2021). Although Ponatinib preserved ATP-linked respiration and OXPHOS-derived ATP production in human fibroblasts, its effects on maximal respiratory capacity and glycolytic metabolism indicate that the compound is not bioenergetically neutral. Ponatinib reduced maximal OCR and spare respiratory capacity in control cells and lowered glycolysis-linked ATP production in cIV-deficient cells, an inhibitory effect on glycolysis previously reported for Ponatinib (Paech et al, 2018). These effects are particularly concerning in a disease context, as cells with impaired OXPHOS already have limited metabolic flexibility and rely on glycolysis to compensate for their respiratory defect, counterbalancing any potential benefit associated with enhanced mitochondrial clearance.

Despite their distinct mechanisms of action and bioenergetic profiles, neither Meptazinol nor Ponatinib improved mitochondrial respiratory function in the respective OXPHOS-deficient models. Because newly generated mitochondria retain the underlying genetic deficiency, increased mitochondrial turnover may not necessarily restore the respiratory capacity of the mitochondrial population. Furthermore, the functional consequences of mitophagy induction depend on its coordination with mitochondrial biogenesis. These findings suggest that effective therapeutic strategies targeting mitochondrial quality control may require coordinated enhancement of mitochondrial turnover and biogenesis, or combination with interventions that directly improve respiratory chain function. In summary, we established a robust high content screening pipeline combining TFEB nuclear translocation, mitophagy induction, and mitochondrial health measurements to identify pharmacological modulators of mitochondrial quality control. Our findings demonstrate that pharmacological enhancement of mitophagy can occur through distinct signaling responses and does not necessarily translate into improved mitochondrial bioenergetics in OXPHOS-deficient cells. These observations highlight the importance of integrating mechanistic and functional validation into drug-discovery strategies targeting mitochondrial quality control and establish a foundation for identifying more effective therapeutic approaches for mitochondrial diseases.

## MATERIALS AND METHODS

### Animals

All animal experiments were carried out in accordance with EU Directive 2010/63/EU (PR474/2021), approved by the local ethical committee (OPBA) at VIMM Institute and Italian Ministry of Health (authorization no. 390/2024-PR). Animal work is compliant with ARRIVE guidelines. The animals were maintained in controlled environmental conditions with cycles of 12 hours light/12 hours dark and they were fed with a standard food diet and water ad libitum. Both males and females were used in this study. Mice (4-8 weeks old) were sacrificed by cervical dislocation, and hindlimb muscles (quadriceps and gastrocnemii) were dissected for MAFs isolation.

### Cell culture

Primary fibroblasts from controls and mitochondrial disease patients (NDUFA6^A178P/c.420+784>T^ and SURF1^R264fs*27^) were acquired from the ‘Cell line and DNA Bank of Genetic Movement Disorders and Mitochondrial Diseases’ of the Telethon Network of Genetic Biobanks (grant GTB12001J). Written informed consent was obtained from all participants or their legal guardians at the source Institution, in accordance with the Declaration of Helsinki. Cells were maintained in EMEM (ATCC, MANASSAS, VA, USA #30-2003) supplemented with 15% FBS (Thermo Fisher, Waltham, MA, USA #A5256701) and antibiotic–antimycotic (Thermo Fisher, Waltham, MA, USA #15240062). Wildtype and MitoQC mice on a C57Bl/6 background are used for the isolation of mouse adult fibroblasts. MAFs were isolated from the skeletal muscle by enzymatic digestion. Mice (4-8 weeks old) were sacrificed by cervical dislocation, and hindlimb muscles (quadriceps and gastrocnemii) were dissected. Muscles were minced and incubated in a Collagenase A solution (Merck, Darmstadt, Germany #10103578001), 1% Penicillin/Streptomycin (Thermo Fisher, Waltham, MA, USA #15140122), 1% Amphotericin B (Thermo Fisher, Waltham, MA, USA #15290026) at 37°C for 30 minutes on a rotating platform. Then, tissue was centrifuged 500 x g for 5 minutes and resuspended in a 1 mg/ml Collagenase/Dispase solution (Merck, Darmstadt, Germany #10269638001), followed by incubation at 37° C for 40 minutes and a second incubation at 37°C for 30 minutes in a 1 mg/mL DNase I solution (Merck, Darmstadt, Germany #10104159001). Muscle homogenate was sequentially filtered and centrifuged at 1200 x g for 10 minutes at room temperature. The pellet was resuspended in mouse fibroblast growth medium (DMEM Glutamax High Glucose, Thermo Fisher, Waltham, MA, USA #31966047, supplemented with 10% FBS and 1% Penicillin/Streptomycin) and seeded in 100 mm dishes. MAFs immortalization was achieved by transfecting the pMSE-SV40LT plasmid encoding for SV40 large T antigen (Schuermann, 1990).

Lentiviral infection of pWPXLd-TFEB-GFP and pLenti-mito-QC (produced by Welgen, Worcester, MA, USA) was performed using 8 mg/ml polybrene (Merck, Darmstadt, Germany #TR-1003) in growth media for 30 h and positive selection was performed by 1 μg/ml puromycin (InvivoGen, San Diego, CA, USA #ant-pr-1). All cell lines were cultured at 37°C and in a controlled atmosphere with 5% humified CO2.

### Treatments

For the treatment of MAFs, DMSO (Merck, Darmstadt, Germany #D2650), Torin 1 (Merck, Darmstadt, Germany #475991), rapamycin (LC Labs #R5000), curcumin C1 (Selleckchem, Houston, TX, USA #S6769), Urolithin A (Merck, Darmstadt, Germany #SML1791), FCCP (Merck, Darmstadt, Germany #C2920), Oligomycin (Merck, Darmstadt, Germany #495455) and the DiscoveryProbe™ FDA-approved Drug Library 125nM (ApexBio, Houston, TX, USA #L1021) were used. When screening the FDA-approved library, each drug was tested in 4 technical replicates within the same 384-well plate. To allow comparison among independent plates, at least 4 positive (Torin 1) and 4 negative control (DMSO) wells were present in each screening plate. Mouse fibroblasts growth medium was changed to mouse fibroblast treatment medium (DMEM Glutamax High Glucose, supplemented with 1% FBS and 1% Penicillin-Streptomycin) 24 hours before treatment. Drug concentrations and duration of treatments are specified in the legends. When described, cells were fixed in 4% formaldehyde for 20 minutes at room temperature.

For the treatment of patient fibroblasts, medium was changed to human fibroblast treatment medium (Phenol red-free DMEM, Thermo Fisher, Waltham, MA, USA #A1443001, supplemented with 1% FBS, 4.5 g/l Glucose (Merck, Darmstadt, Germany #G7021), 2 mM Glutamine (Thermo Fisher, Waltham, MA, USA #25030081), 1 mM Sodium pyruvate (Thermo Fisher, Waltham, MA, USA #11360070) and 1% Antibiotic–Antimycotic) 24 hours before treatment. All drugs were dissolved as indicated in the datasheets: Ponatinib (Selleckchem, Houston, TX, USA #S1490), Cisplatin (Selleckchem, Houston, TX, USA #S1166), Amodiaquine 2HCl 2H_2_O (Selleckchem, Houston, TX, USA #S4589), Mitomycin C (Selleckchem, Houston, TX, USA #S8146), Afatinib (Selleckchem, Houston, TX, USA #S1011), Topotecan (Selleckchem, Houston, TX, USA #S9321), Alendronate sodium 3H_2_O (Selleckchem, Houston, TX, USA #S1624), Cyproheptadine HCl 1.5H_2_O (Selleckchem, Houston, TX, USA #S4635), Meptazinol HCl (Merck, Darmstadt, Germany #M2824). All treatments were performed for 24 hours.

### Plasmids

The sequence encoding for a tagged TFEB-GFP was excised from the pEGFP-N1-TFEB plasmid (Addgene, Watertown, MA, USA #38119) (Roczniak-Ferguson et al, 2012) and inserted into the pWPXLd plasmid (Addgene, Watertown, MA, USA #12258) for lentiviral expression using the In-Fusion® HD Cloning Kit (Takara #639650). The pWPXLd and the pWPXLd-TFEB-GFP vectors (10 µg) were transfected into HEK293T cells, together with 6.55 µg of packaging plasmid (psPAX2, Addgene, Watertown, MA, USA #12260) and 3.5 µg of envelope plasmid (pMD2.G, Addgene, Watertown, MA, USA #12259) using the FUGENE HD transfection reagent (Promega, Madison, WI, USA #E2311).

### Imaging and analysis of TFEB translocation

Images were acquired using the Zeiss LSM900 confocal microscope coupled with a Zeiss EC Plan-Neofluar 20x/0.5 air objective. DAPI and GFP channels were acquired and analyzed with ImageJ software. Briefly, images were thresholded and particles counted. The percentage of TFEB-positive nuclei over the total number of nuclei was calculated to evaluate TFEB translocation into the nucleus. For high-content confocal imaging, TFEB-GFP MAFs were seeded in 384-well plates (Greiner, Kremsmünster, Austria #781091). After treatment, cells were fixed with 4% formaldehyde and stained with DAPI (Thermo Fisher, Waltham, MA, USA #D21490). Imaging was performed with the Operetta CLS High-Content Analysis System (PerkinElmaer, Waltham, MA, USA), using a 20x air objective. Automated analysis of TFEB was performed with Harmony High-Content Analysis Software (PerkinElmaer, Waltham, MA, USA). First, nuclei (50 µm^2^ to 400 µm^2^) and cells (50 µm^2^ to 1200 µm^2^) were identified and filtered by size. A nucleus was considered GFP-positive when the signal intensity was greater than in the cytosol by 1500 a.u. or greater. TFEB translocation was calculated as the ratio between the number of GFP-positive nuclei and the total number of nuclei.

### MitoQC imaging and analysis

For MitoQC imaging, images were collected with one of the following: a Zeiss LSM900 Confocal system with Airyscan equipped with a Zeiss EC Plan-Neofluar 40x/1.3 objective, Operetta CLS High-Content Analysis System (PerkinElmaer, Waltham, MA, USA), using a 40x water objective or the ImageXpress Micro Confocal high-content imaging system (Molecular Devices, San Jose, CA, USA) with a 40x/1.2 water objective. For Zeiss LSM900, 3D image stacks were acquired at optimal Z-distance and reconstructed as Maximum Z-projections. Images were analyzed using Pixel classification segmentation with the Aivia v.10 software. Mitolysosomes (red vesicles only) were counted on a cell-by-cell basis. For images acquired with Operetta system, image analysis was performed using the Harmony High-Content Analysis Software (PerkinElmaer, Waltham, MA, USA). First, nuclei were thresholded (DAPI) and filtered by size (>40 µm^2^). Then, the GFP and mCherry images were merged, and mitochondria were identified using a linear classifier, trained to distinguish between GFP-positive mitochondria and mCherry-positive mitolysosomes based on signal intensity and particle size. The number of mitolysosomes per cell was counted. Mitochondria fragmentation was also analyzed by filtering area (<2 µm^2^) and roundness (>0.85). Cells were classified as “mito-fragmented” when more than 35% of its mitochondria were fragmented. For images acquired with the ImageXpress system, maximum intensity projections of Z-stacks were used for image analysis performed with the MetaXpress software. A TopHat filter was applied on both GFP and mCherry channels. The images were thresholded and transformed into a binary segmentation. This segmented area was used to separate the red vesicles only and count on a cell-by-cell basis.

For the quantification of mitophagy flux, co-treatment for 24 hours with the inhibitors of lysosomal proteases Leupeptin (Merck, Darmstadt, Germany #L2884) and Pepstatin A (Merck, Darmstadt, Germany #P5318) (10 µM each) was performed. Mitophagy flux was calculated as the number of mitolysosomes in presence of Leupeptin and Pepstatin, divided by the mitolysosomes counted in absence of the inhibitors.

### Endogenous TFEB Immunofluorescence and analysis

Following treatment, cells were washed and fixed with 4% paraformaldehyde (Thermo Fisher, Waltham, MA, USA Scientific, Waltham, MA, USA, 043368-9M) in phosphate-buffered saline (PBS; Gibco, Grand Island, NY, USA, Grand Island, NY, USA, 14190-144) at room temperature for 20 min. Following fixation, cells were permeabilized (0.1% Triton X-100 (Sigma-Aldrich, T8787), 0.05% sodium deoxycholate (Sigma-Aldrich, 30970) in PBS) and blocked (5% donkey serum (Millipore Sigma, S30-M) in PBS). Cells were stained with TFEB primary antibody (Cell Signaling, 4240) in blocking solution overnight. After primary antibody removal, a rabbit anti-mouse IgG 647 (Invitrogen, A21239) was used as the secondary antibody and DAPI (Thermo Scientific, D21490) was used to stain nuclei. Between each step, cells were washed with 4x with PBS. Image acquisition was performed using the Image Xpress Micro Confocal high-content imaging system (Molecular Devices, San Jose, CA, USA) with a 20x/0.95 water objective. Z-stacks were acquired, with 1µm between slices (7 slices in total). The maximum intensity projections of Z-stacks were used for image analysis performed with the MetaXpress software. Detection of nuclei was performed on DAPI channel and transformed into a binary segmentation. The segmented nuclei were used to measure the average fluorescence intensity of TFEB. A threshold for average intensity was set equally between conditions to count the TFEB-positive nuclei (% of total nuclei).

### Western Blotting

Cells were pelleted and resuspended in 100 µl of freshly made ice-cold RIPA buffer containing MgCl_2_ 10 mM, NaF 10 mM, Na_3_VO_4_ 1 mM, 1% n-Dodecyl beta-D-maltoside, protease inhibitor 100x (Thermo Fisher, Waltham, MA, USA Scientific, Waltham, MA, USA, 78430), phosphatase inhibitor 20x (Thermo Fisher, Waltham, MA, USA Scientific, Waltham, MA, USA, A32957), then kept on ice for 10 minutes. The extract was centrifuged at 14000rpm in a table-top centrifuge at 4° C for 5 minutes. Protein concentration was determined using the Pierce BCA Protein Assay Kit (Thermo Fisher, Waltham, MA, USA, 23223). Protein samples were mixed with protein loading buffer 4x and 10 % β-mercaptoethanol, then loaded in NuPAGE™ Bis-Tris 4-12% precast polyacrylamide gels (Thermo Fisher, Waltham, MA, USA, NP0321). Gel electrophoresis was performed in a Mini Gel Tank (Thermo Fisher, Waltham, MA, USA) for 75 minutes using a constant voltage of 120 V in MOPS buffer. Proteins were transferred to a methanol-activated Polyvinylidene Fluoride membrane (Thermo Fisher, Waltham, MA, USA Scientific, Waltham, MA, USA, 88520), using a Mini Trans-Blot cell (Bio-Rad, Hercules, CA, USA) at 100 V for 75 min on ice. Next, the blotted membrane was blocked in blocking buffer (3% Bovine serum albumin (BSA; Thermo Fisher, Waltham, MA, USA Scientific, Waltham, MA, USA, BP9703100) in Tris buffered solution containing 0.1% Tween 20 (TBS-T)) for 60 minutes on a shaking platform at RT. The primary antibodies (Phospho-S6 Ribosomal Protein Ser240/244, Cell Signaling, 2215, dilution 1:1000; S6 Ribosomal Protein, Cell Signaling, 2217, dilution 1:1000; vinculin, Sigma, V9131, dilution 1:5000) were incubated overnight at 4° C. Following three washes in TBS-T, the membrane was incubated with HRP-conjugated anti-mouse (Cell Signaling, 7076, dilution 1:3000) or anti-rabbit (Cell Signaling, 7074, dilution 1:3000) IgG HRP-linked antibodies for 1 hour at 4° C. The membrane was once more washed three times with TBS-t after incubation with the secondary antibody. The antibody signal was imaged in a ChemiDoc Imaging System (Bio-Rad, Hercules, CA, USA). Images were analyzed using the Image Lab software (Bio-Rad, Hercules, CA, USA) to quantify band density.

### Lysosomes staining and analysis

Cells were incubated with 50 nM MitoTracker Deep Red FM (Invitrogen, M22426), 50nM LysoTracker Green DND-26 (Molecular Probes, Eugene, OR, USA, L7526), and 1 mg/mL Hoechst (Invitrogen, H3570) in treatment media for 1 hour at 37°C. Cells were washed with warm media and imaged using the Image Xpress Micro Confocal high-content imaging system (Molecular Devices, San Jose, CA, USA) with a 40x-1.2 NA water objective. Z-stacks were acquired, with 0.5 µm between slices (7 slices in total). The maximum intensity projections of Z-stacks were used for image analysis performed with the MetaXpress software. A Gaussian filter was applied on MitoTracker channel for the detection of cytoplasm. Cell-by-cell segmentation was performed by nuclei (Hoechst channel) detection and gaussian filtered cytoplasm (MitoTracker). A binary mask was created for the whole cell containing nuclei added with cytoplasm staining. A TopHat filter of 10 was used for better detection of lysosomes that were detected by using the “finding round object” module keeping the same threshold for all conditions. Total number of lysosomes per cell was averaged and normalized per DMSO treated condition.

### Mitochondrial Membrane Potential

For mitochondrial membrane potential kinetics, cells were incubated with TMRM 10 nM (Thermo Fisher, Waltham, MA, USA #T668), cyclosporine H 1 µM (Merck, Darmstadt, Germany #SML1575) and Hoechst (Thermo Fisher, Waltham, MA, USA) in HBSS (Thermo Fisher, Waltham, MA, USA #14025092) supplemented with 1 g/L of glucose (Thermo Fisher, Waltham, MA, USA #H21486) for 30 minutes. Hoechst (exc. 360-400nm/em. 410-480nm) and TMRM (exc. 520-550nm/em. 560-630nm) were acquired sequentially with a 20x air objective at the Operetta CLS High Content Screening Platform (Perkin Elmer). ΔΨm kinetics was measured by 2 image acquisitions before addition of Oligomycin 1 µM, 12 acquisitions before FCCP 5µM, and 3 acquisitions after. Image analysis was performed using the Harmony 4.8 Analysis software (Perkin Elmer) as described in (Vianello et al, 2023).

For mitochondrial membrane potential and quantification of small, and small-depolarized mitochondria, cells were incubated with TMRE 15 nM (Thermo Fisher, Waltham, MA, USA #T669), MitoTracker Green FM 200 nM (Thermo Fisher, Waltham, MA, USA #M7514) and 1 μg/ml Hoechst (Thermo Fisher, Waltham, MA, USA) for 1 hour at 37°C with 5% CO_2_. FCCP 5 µM was added as positive control. Imaging was performed with the Image Xpress Micro High-Content Imaging System with a 40x water objective. Z-stacks were acquired, with 0.5 µm between slices (6 slices in total). Image analysis was performed using MetaXpress software. Stacks from the TMRE channel were summed for total fluorescence intensity accumulation, while MTG maximum intensity projection was filtered and thresholded to create a mask to measure the average intensity of TMRE signal in the MTG-positive area only. Mitochondria with area <1 µm^2^ were filtered and counted as small mitochondria. The intensity of TMRE channel in the presence of FCCP was used to set up cut-off intensity value for the depolarized mitochondria.

### Intact cell respirometry

Patient-derived skin fibroblasts were seeded at 10000 cells/well (control fibroblasts), 11000 cells/well (cI-deficient fibroblasts), or 15000 cells/well (cIV-deficient fibroblasts) in a Seahorse XFe96/XF Pro Cell Culture Plate (Agilent, Santa Clara, CA, USA). After 48 hours, media was replaced with DMEM supplemented with 4.5 g/l glucose, 1% FBS, 2 mM glutamine, 1 mM sodium pyruvate and antibiotic/antimycotic. On the day before the assay, cells were treated with mitophagy activators for 24 hours. Before the assay was performed, medium was changed to assay medium (DMEM pH 7.4, 4.5 g/l glucose, 5 mM HEPES, 2 mM glutamine and 1 mM sodium pyruvate). Oxygen Consumption Rate (OCR) and Extracellular Acidification Rate (ECAR) were measured in basal conditions and after the sequential addition of 2 µM Oligomycin, 1.5 µM FCCP and 2 µM Antimycin (Merck, Darmstadt, Germany #A8674) + 1 µM Rotenone (Merck, Darmstadt, Germany #557368). After completion of the assay, cells were stained with Hoechst (1 μg/ml in PBS) and nuclei quantified with the Operetta HCS Imaging System. The data was analyzed using the Wave software (Agilent, Santa Clara, CA, USA). OCR and ECAR were normalized to nuclei count per well. All respiratory parameters were normalized by subtracting OCR measured upon injection of Antimycin + Rotenone, representing non-mitochondrial respiration, and calculated using standard protocols (Divakaruni et al, 2014). ATP production rates were then calculated as reported in (Desousa et al, 2023).

### High-Resolution Respirometry

Patient-derived skin fibroblasts were seeded at 180000 cells/flask (control fibroblasts), 220000 cells/flask (cI-deficient fibroblasts), or 250000 cells/flask (cIV-deficient fibroblasts). The next day, cells were treated with mitophagy activators for 72 hours. On the day of assay, cells were pelleted, counted and resuspended in mitochondrial respiration medium MiR05 (0.5 mM EGTA, 3 mM MgCl_2_, 60 mM Lactobionic acid, 20 mM Taurine, 10 mM KH_2_PO_4_, 20 mM HEPES pH 7.1, 110 mM D-sucrose, 1 g/l fatty acid-free BSA), before loading them in the 0.5 ml O2k chambers. Oxygen flux was measured using the NextGen-O2k (Oroboros Instruments, Innsbruck, Austria) and recorded using DatLab 8.0 (Oroboros Instruments, Innsbruck, Austria). The O_2_ flux over time was monitored by polarographic oxygen sensors, which were calibrated daily at experimental temperature of 37 °C. Fibroblasts were permeabilized with subsequent additions of Digitonin. Respiration was then fueled by sequential addition of different substrates: NAD-linked substrates pyruvate, malate and glutamate to induce non-phosphorylating respiration (leak respiration); ADP to induce c-I respiration coupled to ATP synthesis (basal respiration cI); succinate to induce cII-linked respiration, providing the additive effect of donating electrons to the Q-junction from both cI and cII (basal respiration cI+cII). The uncoupler CCCP was added to induce maximum electron transfer capacity (maximal respiration cI+cII), and afterwards cI was blocked with rotenone to measure maximum electron transfer capacity from cII (maximal respiration cII). All values were normalized to oxygen flux measured in presence of antimycin A. The maximum electron transfer capacity of cIV (maximal respiration cIV) was measured upon addition of ascorbate and tetramethylphenylenediamine and normalized on oxygen flux in presence of sodium azide.

### Statistical analysis

Results are expressed as “mean ± standard error of the mean (SEM)”. Prism 8.0.1 (GraphPad, Boston, MA, USA) was used to perform all statistical analyses and to create all graphs. Unpaired Student’s *t* test was used to compare two groups. One-way analysis of variance (ANOVA) with Dunnett’s post hoc test was used to compare multiple groups. The level of statistical significance is set at p-value≤0.05. Exact p-values and number biological replicates are reported in the figure legends. Individual dots in the graphs represent technical or biological replicates, as stated in the figure legend. The analysis of TFEB screening was performed using Microsoft Excel and RStudio (packages DescTools, multcomp, ggplot2). Quality control was performed by calculating the plate Z’-factor (Zhang et al, 1999), and plates with Z’-factor<0.5 were excluded from further analyses. One-way ANOVA followed by a post-hoc Dunnett’s HSD was performed for TFEB translocation in each screening plate independently. Cell counts after treatment were normalized to the negative control DMSO, and the drugs reducing cell count by >50% were excluded from the next screening step. Enrichment analysis of the drug pathways, as annotated in the library datasheet, was performed by Fisher’s exact test between significant and non-significant TFEB activators.

For analysis of MitoQC, one-way ANOVA followed by post-hoc Tukey HSD test was used for statistical analysis on mitolysosome number per cell and mito-fragmented cells in each plate independently. Drugs causing mitochondrial fragmentation were excluded from the list of mitophagy activators when not significantly different from FCCP.

## ACKNOWLEDGMENTS

This work was supported by; AFM-Téléthon (23706 to C.V. and 24962 to E.F.-V.); Telethon Foundation (GGP20013 and GSP24003 to C.V. and GJC21014 to E.F.-V.), Associazione Luigi Comini Onlus (MitoFight2, to CV), The Lily and POLG Foundation Awards (to CV); PNRR Mission 4, Component 2, Investment 1.4-CN00000041 Spoke 1 funded by the European Union – NextGenerationEU, and MC_PC_21046 to establish a National Mouse Genetics Network Mitochondria Cluster (MitoCluster) (to CV).

## AUTHOR CONTRIBUTIONS

C.B. and C.V. conceived and supervised the study. G.G., V.B., A.E., S.C., L.F.-D.-R., N.S., and D.S. performed experiments and analyzed data. M.G. designed the screen. M.S., E.F.-V., and O.S. provided resources and supervision. C.V., O.S., and E.F.-V. acquired funding. G.G., L.F.-D.-R., C.B., and C.V. wrote the manuscript. All authors reviewed and approved the final version.

## ADDITIONAL INFORMATION

The authors declare no competing interests.

## AVAILABILITY OF DATA AND MATERIALS

The data that support the findings of this study are available from the corresponding authors.

## USE OF AI TOOLS

Claude has been used to improve the language. The authors carefully checked the AI manipulated text.

